# Optimizing FreeSurfer’s Surface Reconstruction Parameters for Anatomical Feature Estimation

**DOI:** 10.1101/2023.01.02.522457

**Authors:** Zvi Baratz, Yaniv Assaf

## Abstract

Magnetic resonance imaging (MRI) is a powerful tool for non-invasive imaging of the human body. However, the quality and reliability of MRI data can be influenced by various factors, such as hardware and software configurations, image acquisition protocols, and preprocessing techniques. In recent years, the introduction of large-scale neuroimaging datasets has taken an increasingly prominent role in neuroscientific research. The advent of publicly available and standardized repositories has enabled researchers to combine data from multiple sources to explore a wide range of scientific inquiries. This increase in scale allows the study of phenomena with smaller effect sizes over a more diverse sample and with greater statistical power.

Other than the variability inherent to the acquisition of the data across sites, preprocessing and feature generation steps implemented in different labs introduce an additional layer of variability which may influence consecutive statistical procedures. In this study, we show that differences in the configuration of surface reconstruction from anatomical MRI using FreeSurfer results in considerable changes to the estimated anatomical features. In addition, we demonstrate the effect these differences have on within-subject similarity and the performance of basic prediction tasks based on the derived anatomical features.

Our results show that although FreeSurfer may be provided with either a T2w or a FLAIR scan for the same purpose of improving pial surface estimation (relative to based on the mandatory T1w scan alone), the two configurations have a distinctly different effect. In addition, our findings indicate that the similarity of within-subject scans and performance of a range of models for the prediction of sex and age are significantly effected, they are not significantly improved by either of the enhanced configurations. These results demonstrate the large extent to which elementary and sparsely reported differences in preprocessing workflow configurations influence the derived brain features.

The results of this study are meant to underline the importance of optimizing preprocessing procedures based on experimental results prior to their distribution and consecutive standardization and harmonization efforts across public datasets. In addition, preprocessing configurations should be carefully reported and included in any following analytical workflows, to account for any variation originating from such differences. Finally, other representations of the raw data should be explored and studied to provide a more robust framework for data aggregation and sharing.

## 1. Introduction

Multi-center studies and data aggregation initiatives have become a standard resource in the field of neuroimaging and a clear prerequisite for the long-term advancement of the field (Horien et al., 2021; Laird, 2021; Van Horn & Toga, 2009). The creation and utility of such datasets introduces numerous technical and methodological challenges which have become an increasingly popular topic of research, and often debate. However, even if all data is archived, preprocessed, and distributed in a completely tested, reproducible, and documented manner, working with repositories of datasets at a large scale requires compromises to be made. The variability and complexity of MRI acquisition and preprocessing predetermines highly heterogeneous results (Borghi & Gulick, 2018; Botvinik-Nezer et al., 2020; Gronenschild et al., 2012), and it is practically impossible to standardize acquisition protocols and preprocessing procedures across all research centers and application domains. Routine procedures, such as providing access to raw data, modifying analysis configurations and integrating new workflows, all become significantly more difficult to support with every increase in scale.

Assuming variation in the many procedures and configurations applied to raw research data is inherent (and to an extent desirable), it is imperative we continuously refine our understanding of the implications of each decision that is made and the significance of various steps and parameters used in the preparation and distribution of our data. This study focuses on sMRI preprocessing, which is the common denominator of the great majority of MRI-based neuroimaging research, and more specifically on the reliability of cross-sectional anatomical statistics calculated with FreeSurfer’s surface reconstruction pipeline (Fischl, 2012) using a subset of available execution configurations. FreeSurfer is a popular and freely available open-source neuroimaging toolkit for processing, analyzing, and visualizing human brain MR images (https://surfer.nmr.mgh.harvard.edu/). It has previously been demonstrated that FreeSurfer successfully captures subtle morphological changes in brain structure (Lehmann et al., 2010; Salat et al., 2009), and several studies have already pioneered the evaluation of variation originating from differences in FreeSurfer versions, operating system, scan session, head-tilt, inter-scan interval, acquisition sequence, and preprocessing stream (cross-sectional or longitudinal) (Dickerson et al., 2008; Gronenschild et al., 2012; Han et al., 2006; Hedges et al., 2022; Jovicich et al., 2009; Knussmann et al., 2022). Such explorations are key for the sustainability of multi-site collaborations and research leveraging data from heterogeneous sources in the field in general.

Another perspective on the reliability and robustness of FreeSurfer-derived anatomical statistics deals with participant identification and classification. Identification refers to the ability to “recognize” an individual from a particular sample, i.e., the model has the same number of target classes as the number of participants it has been trained on. New samples are associated by the model to the most likely participant. Classification, on the other hand, provides some grouping or categorization in which the number of classes is smaller than the number of subjects, e.g., by age group, sex, dominant hand, etc. (Valizadeh et al., 2018). This article will provide a summary of the results of two types of prediction tasks: (a) distinguishing between pairs of MPRAGE scans belonging to the same subject versus pairs belonging to different subjects, and (b) predicting a subset of basic participant attributes, namely sex, age, and BMI. Each task will be trained on an identical set of scans, but using anatomical statistics generated by different FreeSurfer execution configurations.

The purpose of this study is to expand on previous studies and determine the most reliable and informative configuration to use for the estimation of anatomical brain features using FreeSurfer’s surface reconstruction. For this purpose, a subset of the possible execution configurations will be run and compared in two ways; within-subject variability (calculated as cosine similarity), and prediction performance for sex and age using a sample of commonplace estimators. The hypothesis driving this study is that differences in surface reconstruction parameters have a strong effect on the estimated anatomical features, and that these differences can and should be optimized for the benefit of consecutive statistical procedures. This endeavor is meant to serve as a preliminary framework for following studies to tune preprocessing procedures in a manner which optimizes performance for a given statistical effect.

## 2. Materials and Methods

### 2.1. Participants

This study uses a dataset collected from 121 participants (49 female and 72 male), 20–69 years old (mean: 31.22, SD: 9.03; see figure 1 for participant age and sex distribution). Participants were neurologically and radiologically healthy, with no history of neurological disease, and normal appearance in a clinical MRI protocol. The imaging protocol was approved by the Institutional Review Boards of Sheba Tel HaShomer Medical Center and Tel Aviv University, where the MRI investigations were performed. All participants provided signed informed consent before enrollment in the study.

**Figure 1.**
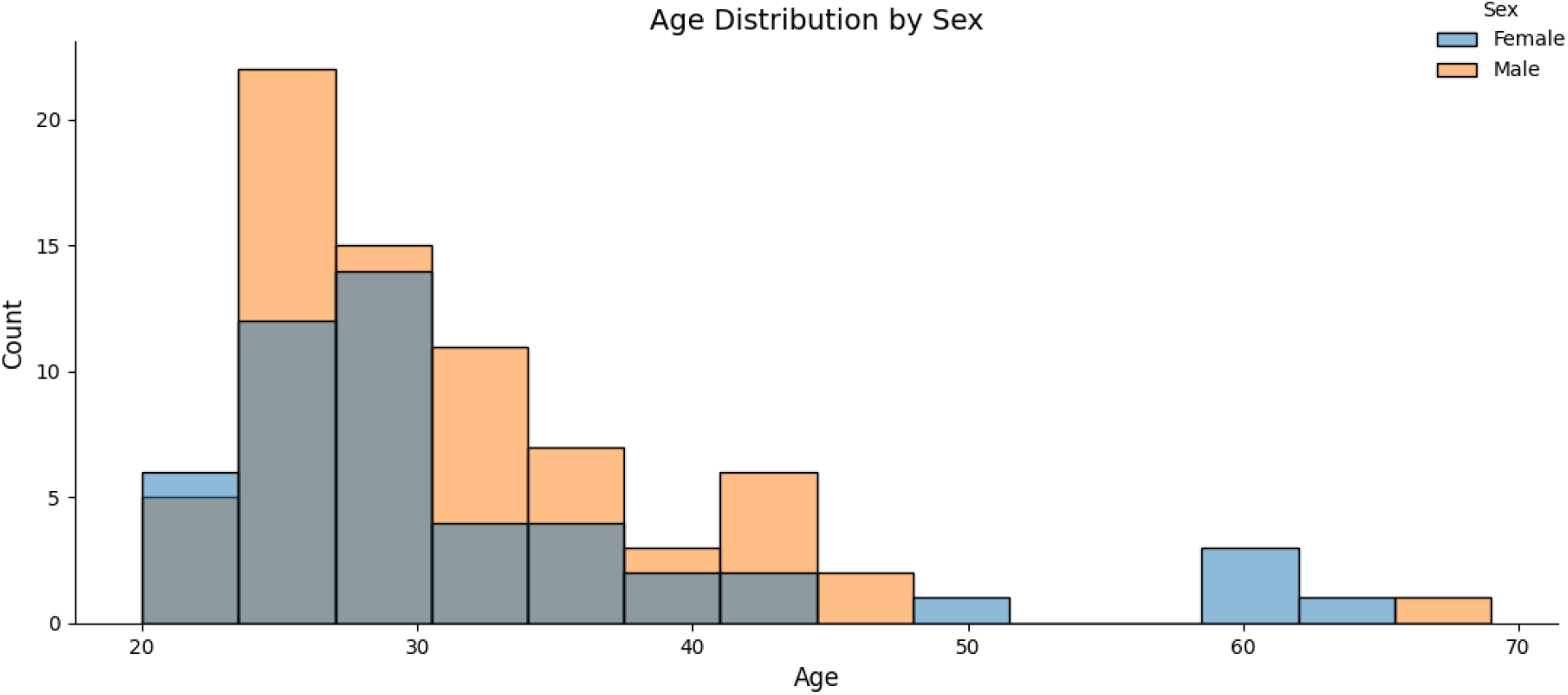
Participant age distribution by sex.

**Figure 2.**
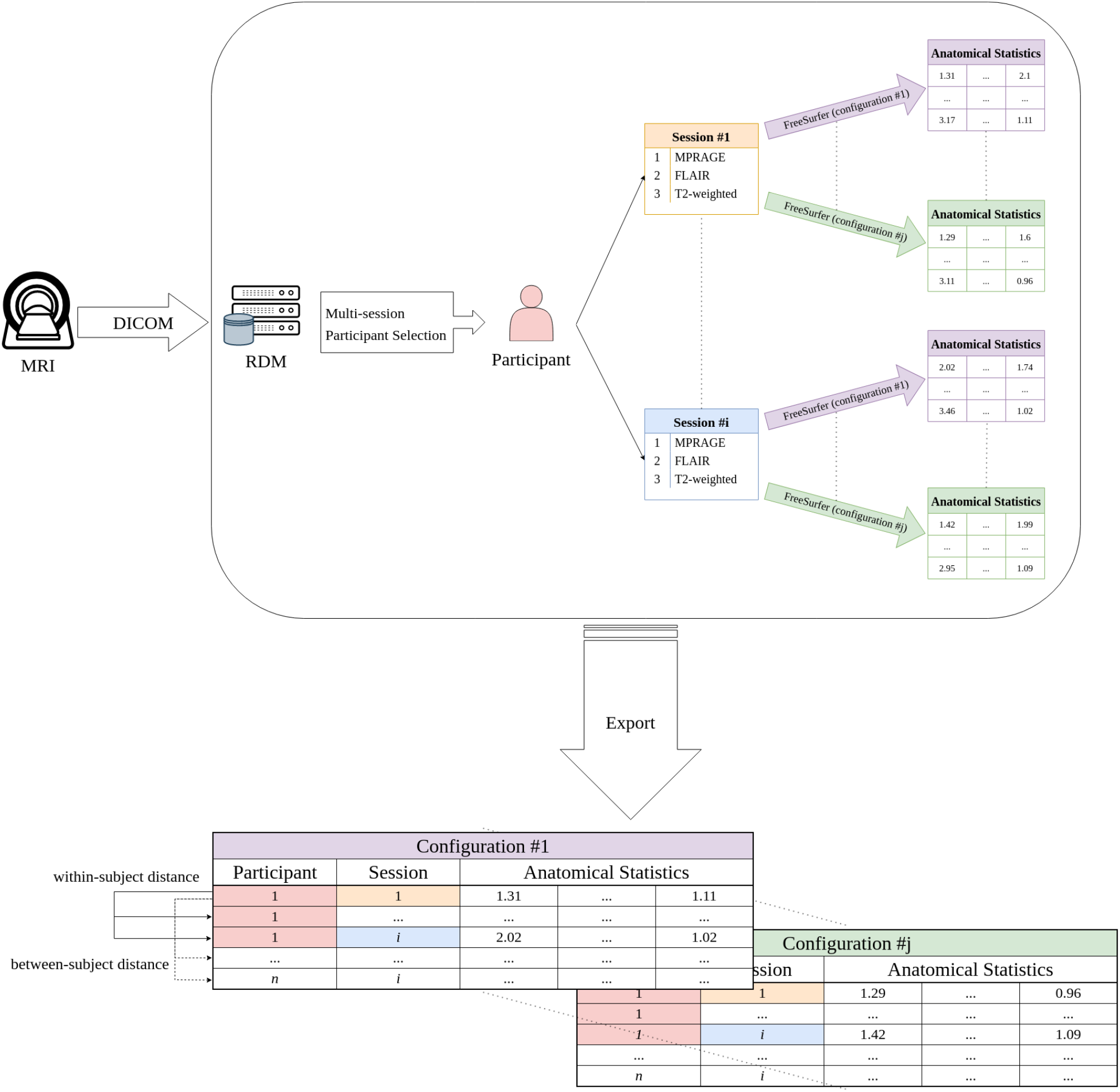
An overview of the anatomical MRI preprocessing workflow. The TAU Brain Bank Initiative’s dataset was filtered for multi-session subjects, and a subset of possible FreeSurfer execution configurations was processed for each scan independently.

### 2.2. MRI Acquisition

Participants were scanned in compliance with the TAU Brain Bank Initiative’s standardized scanning protocol at the Alfredo Federico Strauss Center for Computational Neuroimaging at Tel Aviv University. Scans were acquired with a 3T Siemens MAGNETOM Prisma MRI scanner (Siemens Medical Solutions, Erlangen, Germany) using a 64-channel RF head coil. For the purposes of this study, only the sMRI subset of the protocol which is used for anatomical preprocessing with FreeSurfer will be included (see Table 1 for an overview of acquisition parameters).

**Table 1.** Summary of the TAU Brain Bank Initiative’s sMRl acquisition parameters.

| Sequence | TR (ms) | TE (ms) | TI (ms) | Spatial Resolution (mm<sup>3</sup>) | Number of Slices | Flip Angle | FOV (mm) | Acquisition Matrix | Orientation |
| --- | --- | --- | --- | --- | --- | --- | --- | --- | --- |
| MPRAGE | 2400 | 2.78 | 1000 | 0.9×0.9×0.9 | 176 | 8° | 172 × 230 | 192 × 256 | Axial |
| T2w | 3200 | 554 | - | 0.9×0.9×0.9 | 176 | 120° | 172 × 230 | 192 × 256 | Axial |
| FLAIR | 8000 | 81 | 2370 | 0.72×0.72×4 | 52 | 150° | 173 × 231 | 168 × 320 | Axial |

### 2.3. MRI Data Analysis

FreeSurfer v7.2.0 was used to preprocess the anatomical data using the cross-sectional processing stream (i.e., as individual datasets, rather than longitudinally) with a number of predefined configurations. While FreeSurfer does also offer a longitudinal processing stream (which uses results from the cross-sectional analysis of all time-points to create an unbiased within-subject template), using it would bias the estimation of within-subject similarity across scanning sessions. All runs were executed on a single 64-bit Debian GNU/Linux workstation. Anatomical statistics were extracted for the Destrieux atlas (Fischl et al., 2004), which is the most detailed atlas included with FreeSurfer’s default reconstruction pipeline’s results, and consists of 74 regions per hemisphere.

According to the FreeSurfer documentation site (see https://surfer.nmr.mgh.harvard.edu/fswiki/FreeSurferWiki), using T2w or FLAIR data (mutually exclusive) improves pial surface estimation (see https://surfer.nmr.mgh.harvard.edu/fswiki/recon-all), and specifying *3T* and *mprage* (non-mutually exclusive) alters various analysis assumptions and steps to optimize for the corresponding magnetic field magnitude and acquisition protocol (see https://surfer.nmr.mgh.harvard.edu/fswiki/OtherUsefulFlags). Table 2 provides a summary of the execution configurations included in this study.

**Table 2.** FreeSurfer’s cross-sectional anatomical preprocessing workflow configurations included in this study.

| Use T2 | Use FLAIR | MPRAGE | 3T |
| --- | --- | --- | --- |
| ✗ | ✗ | ✗ | ✗ |
| ✓ | ✗ | ✗ | ✗ |
| ✗ | ✓ | ✗ | ✗ |
| ✓ | ✗ | ✓ | ✗ |
| ✗ | ✓ | ✓ | ✗ |
| ✓ | ✗ | ✓ | ✓ |
| ✗ | ✓ | ✓ | ✓ |

While these options are listed in the documentation website, only a very limited description of their respective effects is provided. In addition, it is unclear to what extent these options are, in fact, utilized by FreeSurfer users, and no existing evaluation of either expected effects or usage metrics could be found at the time of writing.

### 2.4. Statistical Analysis

#### 2.4.1. Comparison of Anatomical Statistics

Region-wise statistics were generated per scan for the detailed execution configurations (see Table 2). Each Destrieux atlas region was represented by four estimated statistics: mean and standard deviation of the cortical thickness, surface area, and gray matter volume. Both whole-brain and region-wise distributions of values were compared.

##### 2.4.1.1. Two-sample Kolmogorov–Smirnov test

The two-sample Kolmogorov–Smirnov (KS) test is a nonparametric test which quantifies the distance between the two samples’ empirical distribution functions and evaluates the probability of them having been sampled from the same probability distribution. The KS test was applied using SciPy (Virtanen et al., 2020) (see https://docs.scipy.org/doc/scipy/reference/generated/scipy.stats.ks_2samp.html) to all combinations of FreeSurfer execution configurations and p-values were corrected to account for multiple comparisons with statsmodels (Seabold & Perktold, 2010) (see https://www.statsmodels.org/stable/generated/statsmodels.stats.multitest.multipletests.html) using the Bonferroni method (Bonferroni, 1935).

#### 2.4.2. Similarity Learning

Within and between-subject distances were calculated as Euclidean, Manhattan, and cosine distances for each FreeSurfer execution configuration. While the Euclidean distance generally provides the most intuitive interpretation of distance in space, Manhattan distance often performs better in high dimensional spaces and is less sensitive to outliers (Aggarwal et al., 2001). In contrast to those, cosine distance is calculated from the angle between the two vectors, which offers the advantage of generating values between 0 and 1 for any dimensionality. All estimated anatomical statistics were standardized within execution configuration before distance calculation.

#### 2.4.3. Model Selection and Optimization

Some commonly used estimators were selected for each prediction task and compared over a predetermined grid of hyperparameters (see figure 14 in the supplementary material). Sex classification was conducted with logistic (logit) regression, as well as random forest and support vector classifiers (RFC and SVC). For age prediction, we used the elastic net and support vector regression (SVR) estimators. All train and test scores are retained in the process of hyperparameter optimization for inspection and evaluation of overall model performance.

## 3. Results

### 3.1. Differences in Estimated Anatomical Statistics

#### 3.1.1. Two-sample Kolmogorov-Smirnov (KS) Test

To first provide a statistical indication that the empirical distributions of the collected metrics differ across execution configurations, a two-sample KS test was applied to compare both whole-brain and region-wise values and estimate the probability of them originating from the same probability distribution as the default configuration’s empirical distribution function (p-values were corrected using the Bonferroni method to account for multiple comparisons). The results indicate highly significant and widespread changes in the mean and standard deviation of the estimated cortical thickness relative to the default across all execution configurations. In addition, FLAIR-complemented results demonstrate a significant difference in the estimated gray matter volume, whereas T2w do not. For a complete overview of the KS test results, see figure 15 and figure 16 included in the Supplementary Material.

#### 3.1.2. Average Cortical Thickness

Average cortical thickness was calculated across all participants for each Destrieux atlas region and FreeSurfer execution configuration. Figure 3 shows the distributions and differences of mean average cortical thickness values per region. While both T2w and FLAIR are considered to improve pial surface estimation, which explains the large effect both have on the estimated cortical thickness, this figure demonstrates an opposite trend manifested by each; adding a T2w reference generally increases the estimated average thickness across nearly all regions, whereas FLAIR results in some regions increasing and some decreasing and an overall reduction. Curiously, using the *MPRAGE* flag has no discernible effect at all.

**Figure 3.**
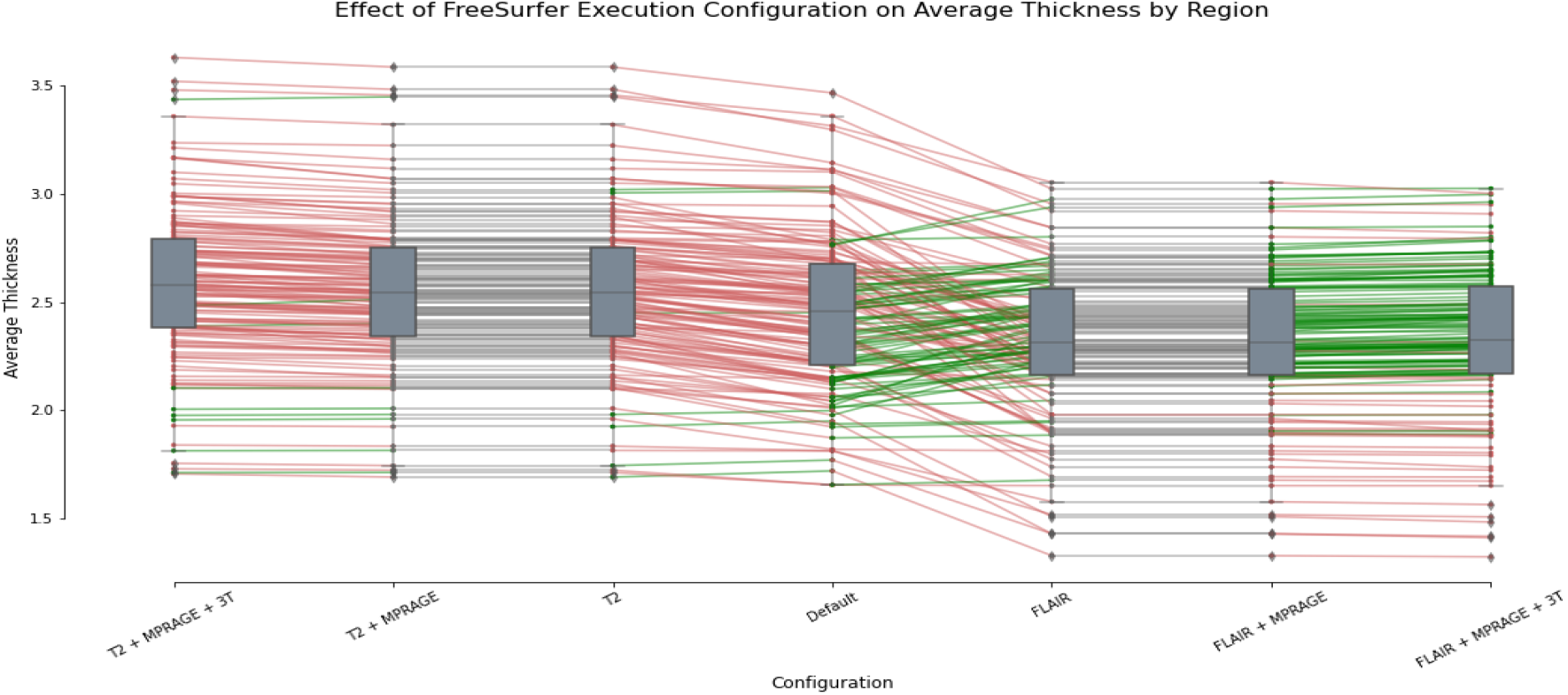
Mean estimated average thickness across participants per execution configuration. Green lines indicate an increase in value and red indicate decrease (from left to right).

The projection of the mean differences onto the standard MNI152 template (see figure 4 and figure 5) reveals that the widespread increase in the estimated cortical thickness when T2w scans are used is particularly evident in the superior frontal gyrus (SFG). In addition, the opposite trends displayed in figure 5 for the results with FLAIR scans reflect a general decrease in the estimated cortical thickness of gyri and increase in sulci. For a complete summary of the estimated anatomical statistics across execution configurations by atlas region, see figure 17 in the Supplementary Material.

**Figure 5.**
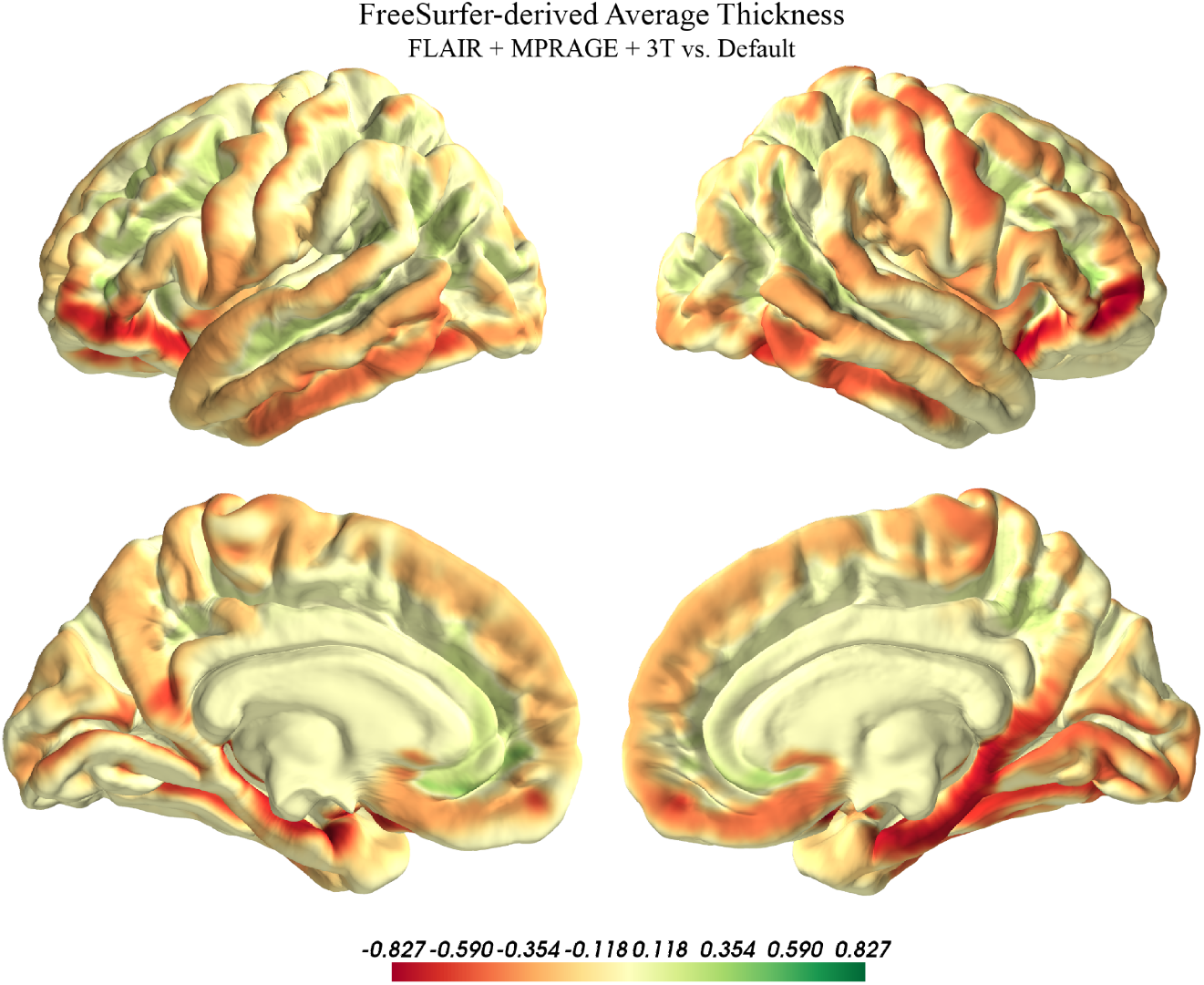
Difference between average cortical thickness estimated with FreeSurfer’s surface reconstruction pipeline including a FLAIR scan versus the default execution configuration. Values were averaged across participants by region and the differences projected onto the MNI152 standard template.

#### 3.1.3. Standard Deviation of Cortical Thickness

Standard deviation of the cortical thickness was calculated across all participants for each Destrieux atlas region and FreeSurfer execution configuration. Figure 6 shows the distributions and differences of values per region. These results demonstrate a general decrease for the T2w execution configuration (contrary to the overall increase in absolute value), with FLAIR again showing opposite trends in different regions.

**Figure 6.**
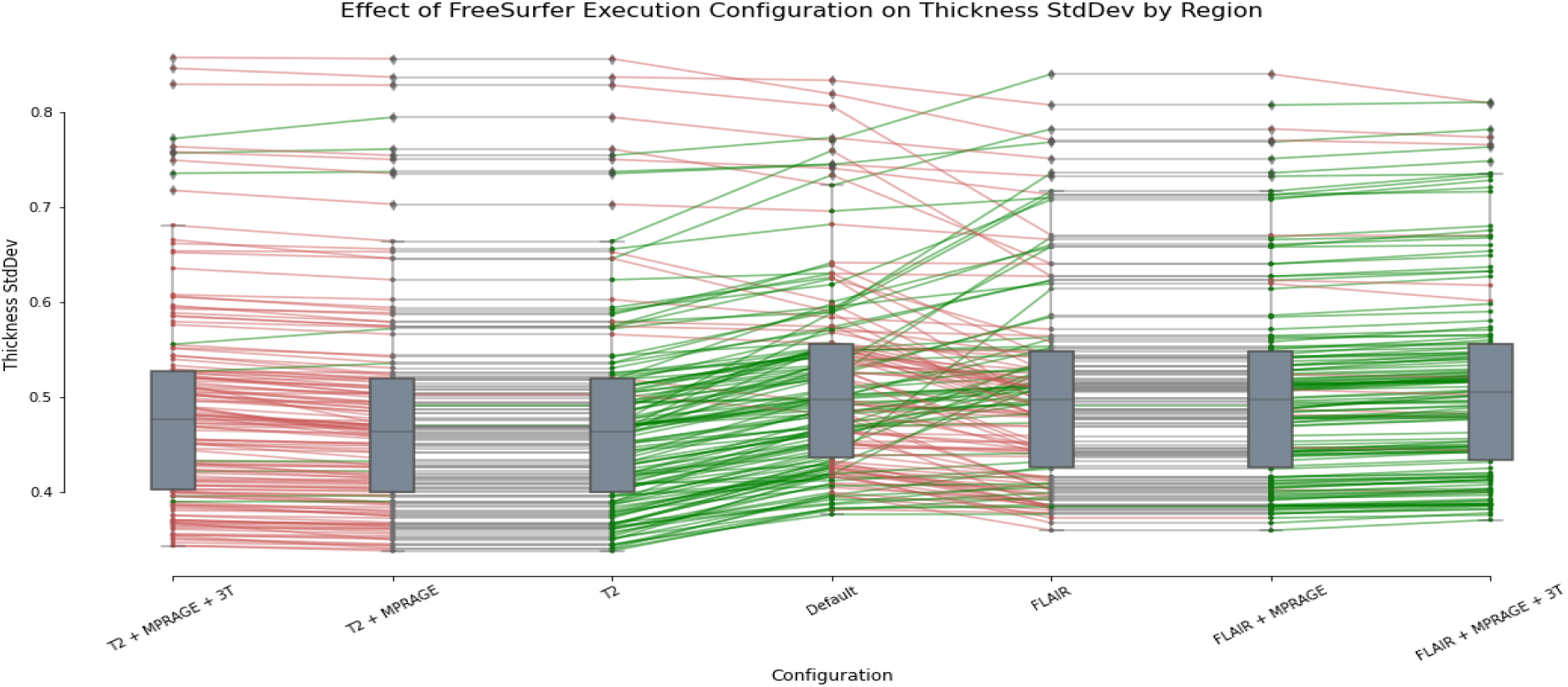
Mean estimated thickness standard deviation across participants per execution configuration. Green lines indicate an increase in value and red indicate decrease (from left to right).

**Figure 7.**
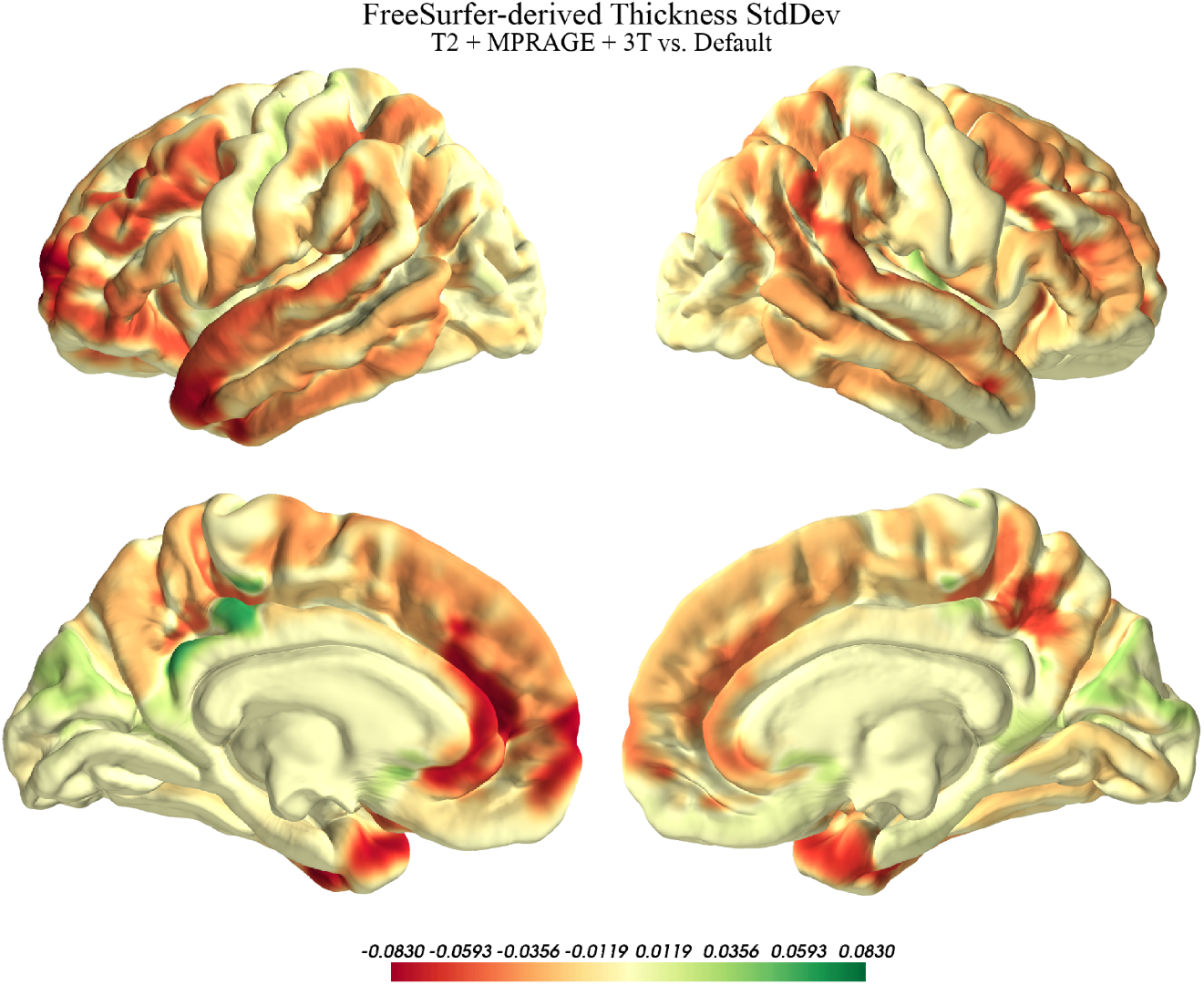
Difference between the standard deviation of the cortical thickness as estimated with FreeSurfer’s surface reconstruction pipeline including a T2w scan versus the default execution configuration. Values were averaged across participants by region and the differences projected onto the MNI152 standard template.

**Figure 8.**
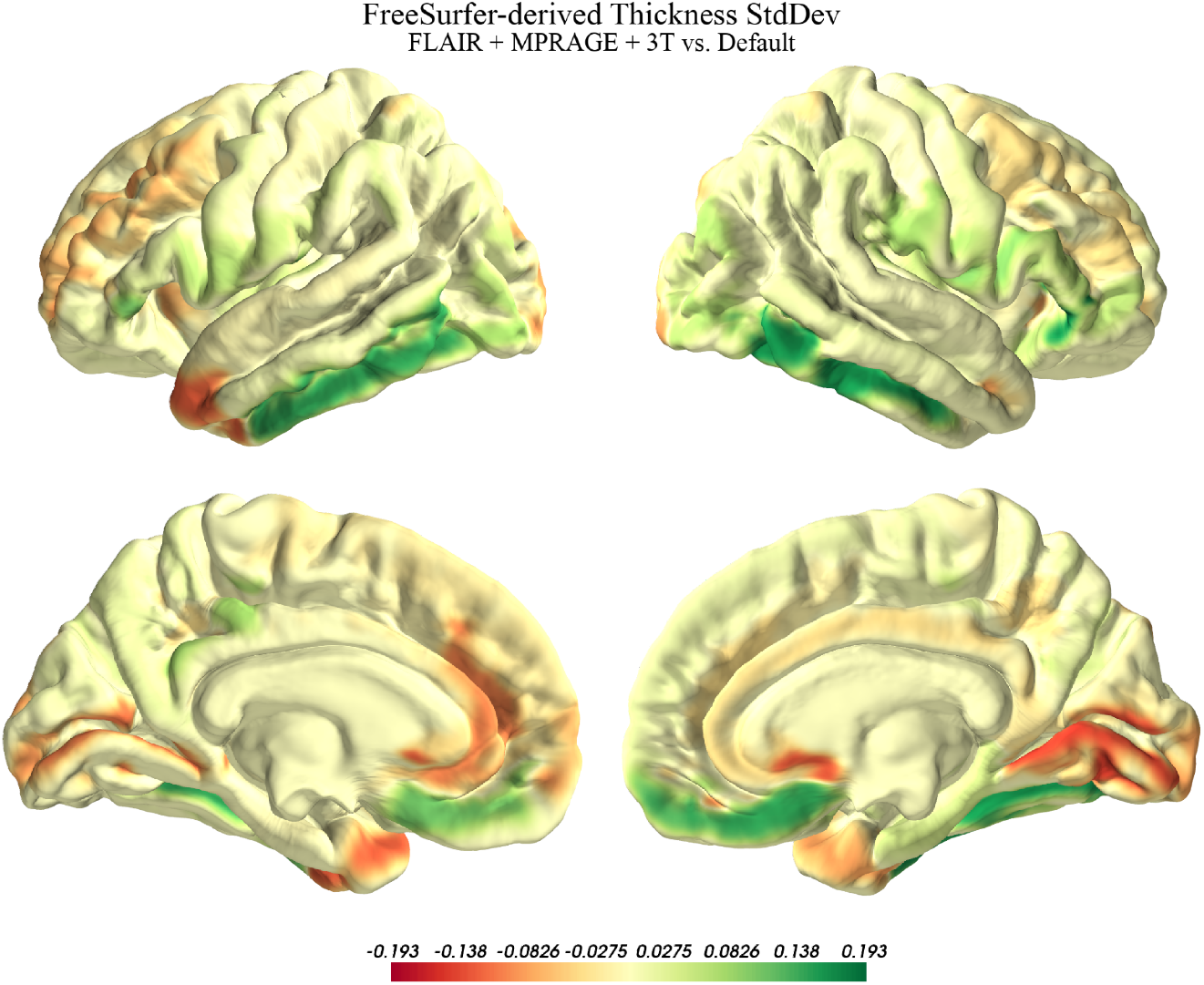
Difference between the standard deviation of the cortical thickness as estimated with FreeSurfer’s surface reconstruction pipeline including a FLAIR scan versus the default execution configuration. Values were averaged across participants by region and the differences projected onto the MNI152 standard template.

#### 3.1.4. Gray Matter Volume

Estimated gray matter volume changed significantly only for FLAIR-complemented analyses, resulting in a general decrease in values and overall variability. Figure 9 shows the distribution of region-wise mean gray matter volume, and Figure 10 shows a projection of the difference between the FLAIR and default execution configurations.

**Figure 9.**
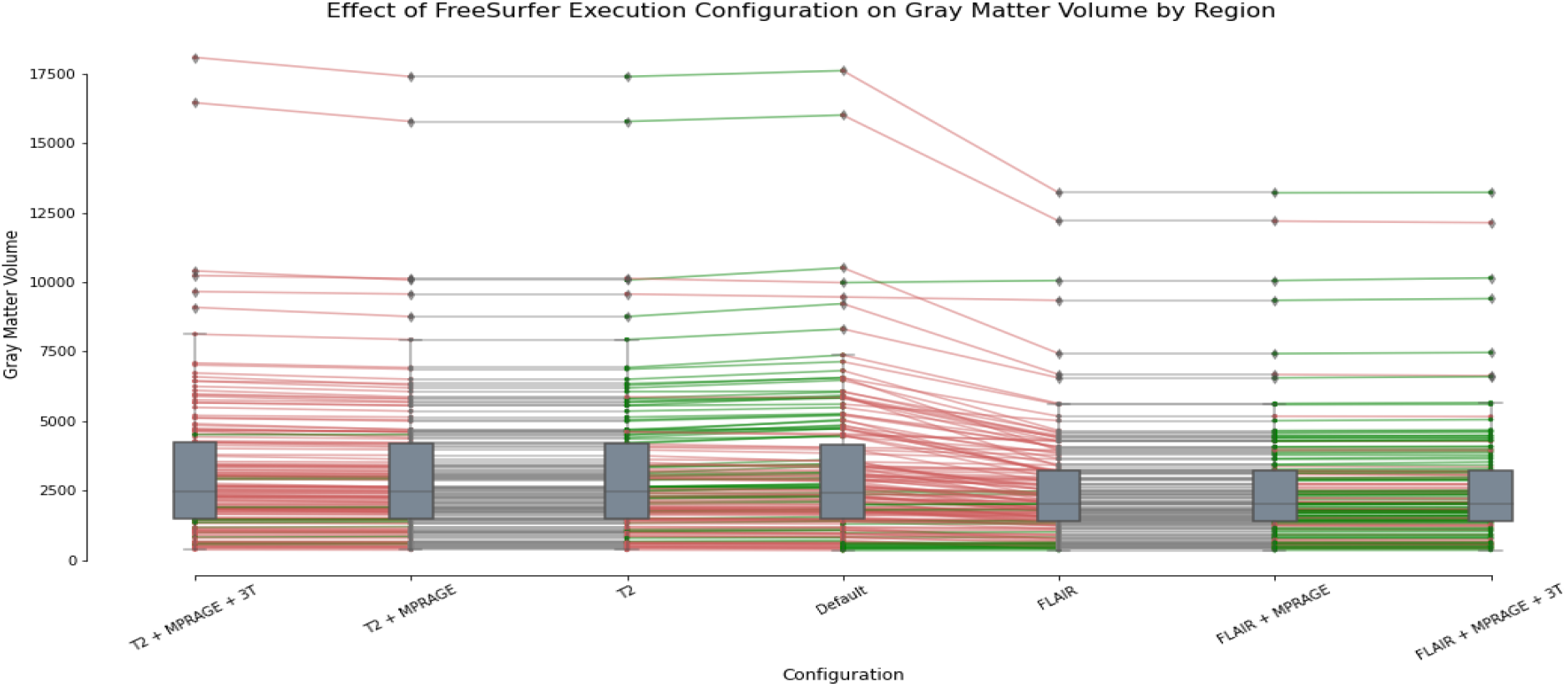
Mean gray matter volume across participants per execution configuration. Green lines indicate an increase in value and red indicate decrease (from left to right).

**Figure 10.**
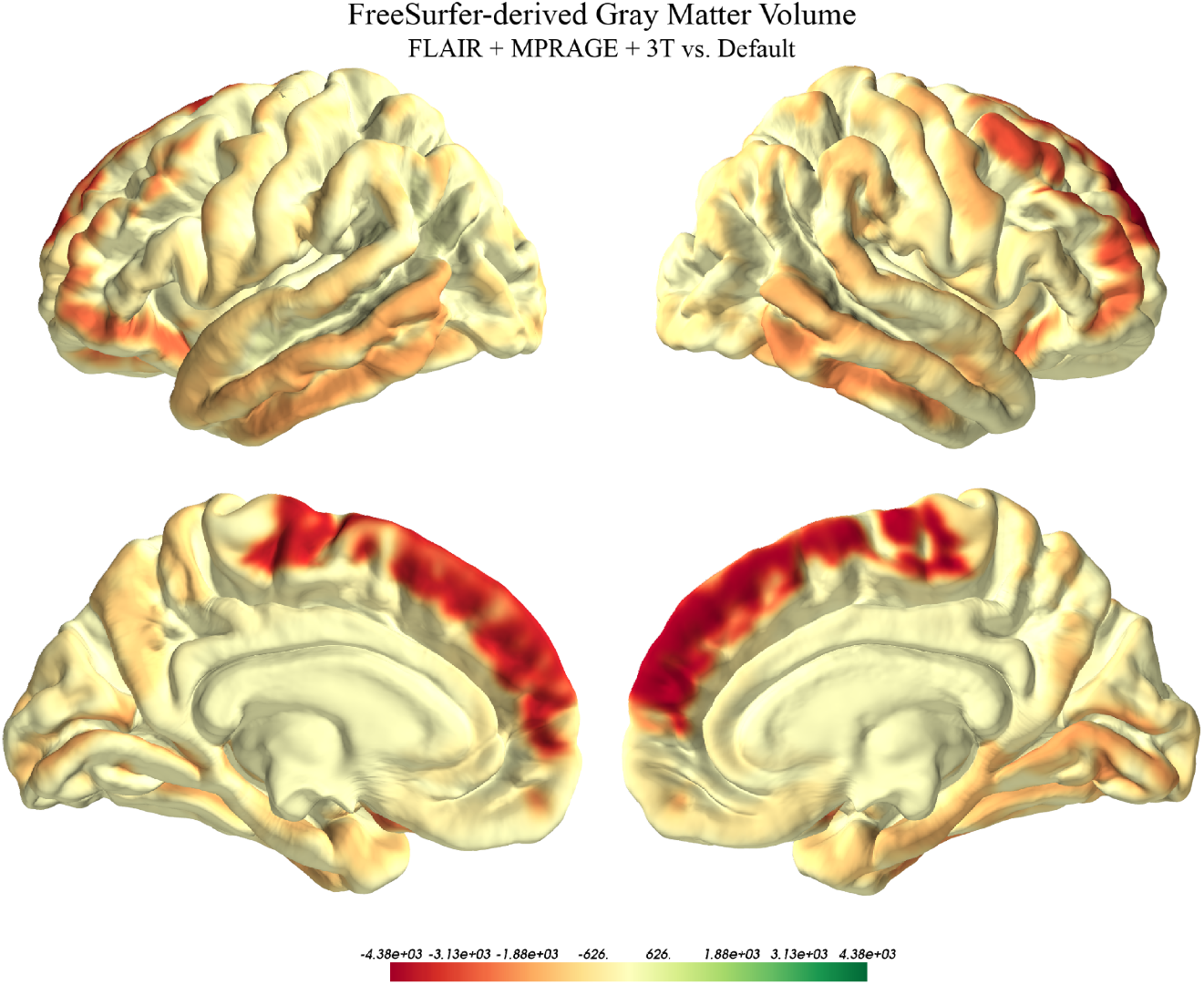
Difference between the gray matter volume estimated with FreeSurfer’s surface reconstruction pipeline including a FLAIR scan versus the default execution configuration. Values were averaged across participants by region and the differences projected onto the MNI152 standard template.

**Figure 11.**
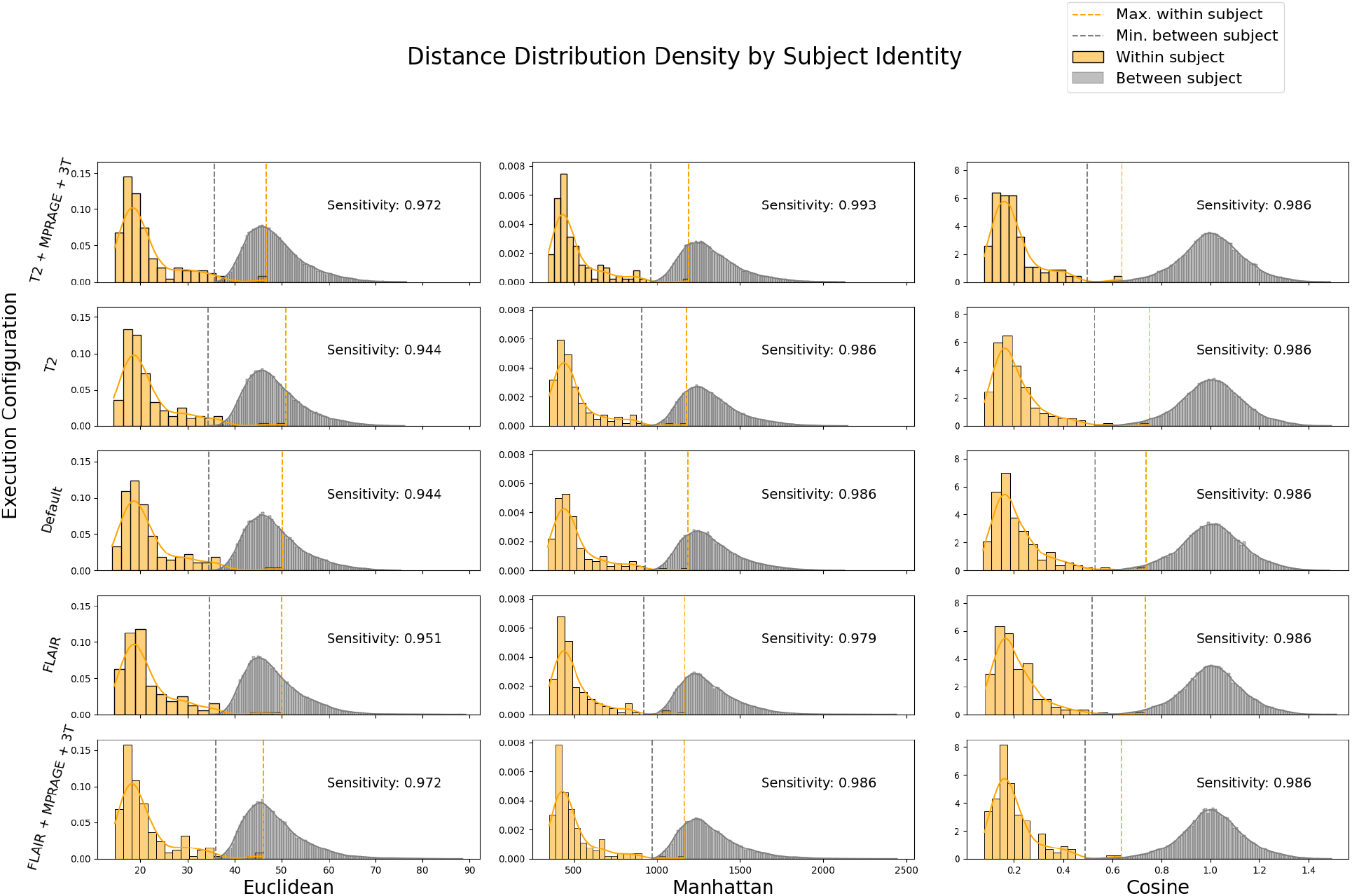
Within vs. between-subject distance metric distributions across FreeSurfer execution configurations.

**Figure 12.**
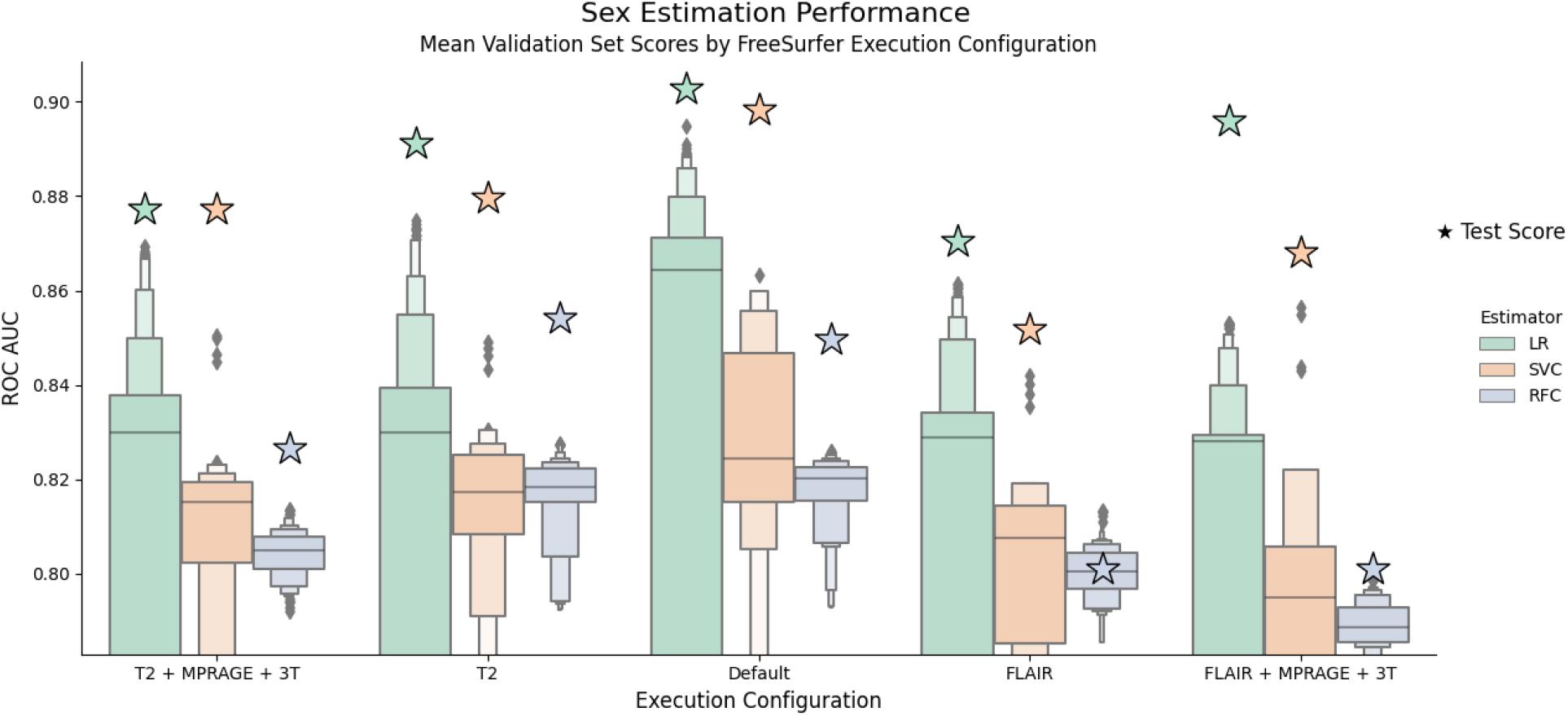
Distribution of mean validation scores (ROC AUC) across LR, SVC, and RFC parameters by FreeSurfer execution configuration for the classification of sex. Stars represent out-of-sample test scores.

**Figure 13.**
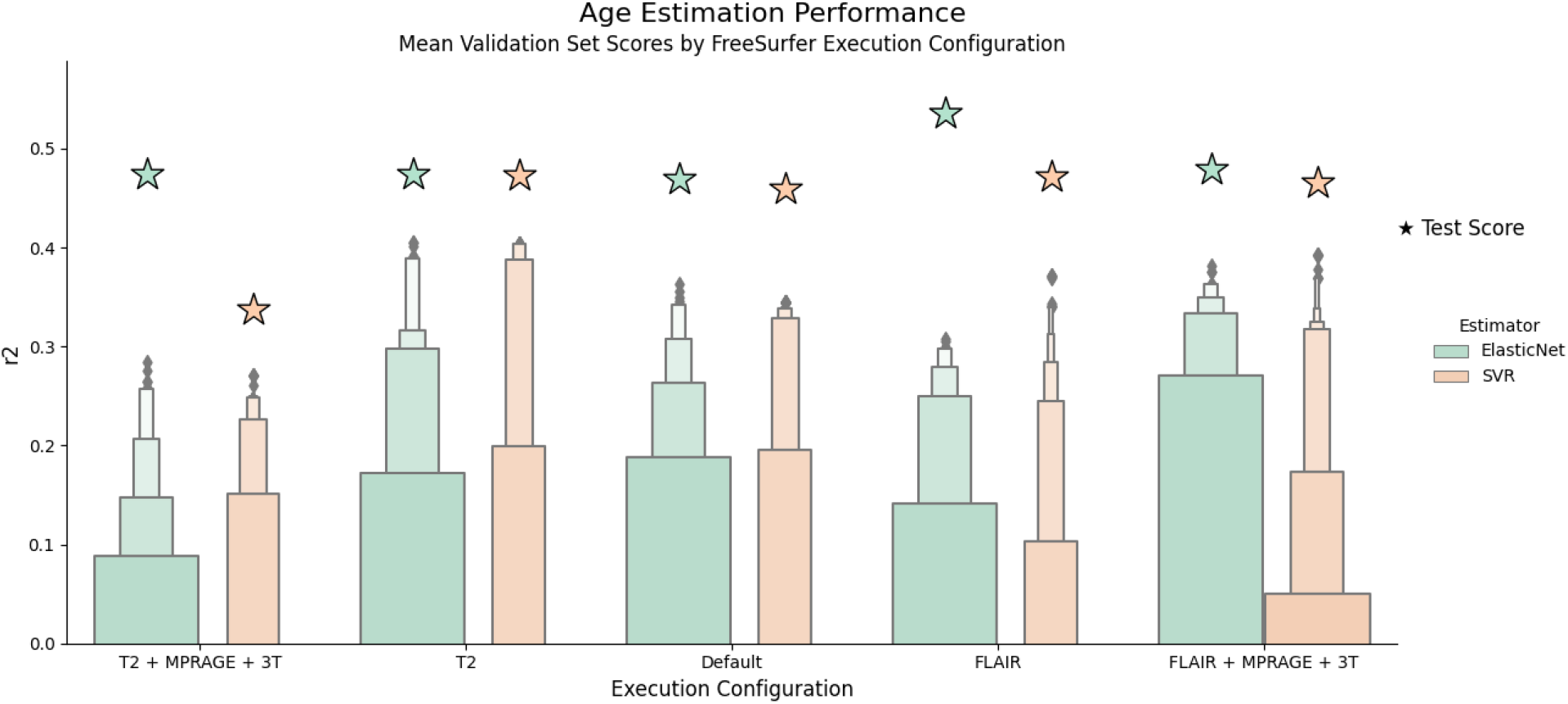
Distribution of mean validation scores (r^2^) across elastic net and SVR parameters by FreeSurfer execution configuration for the classification of sex. Stars represent out-of-sample test scores.

The decrease in estimated gray matter volume when using FLAIR scans to enhance pial surface estimation is most evident in the frontal lobe, and particularly in the SFG.

### 3.2. Similarity Learning

Three distance metrics were calculated and used to evaluate within-subject similarity relative to between subjects. All results were standardized by execution configuration (i.e., all features have a mean close to 0 and standard deviation close to 1 across runs with identical FreeSurfer parameters) and distances were calculated for every possible combination of runs in the dataset.

These results show excellent separation between within and between-subject scans for all calculated distance metrics. Using the minimal between-subject distance as a simple classification threshold provides within-subject sensitivity scores ranging from 94.4% to 99.3% (with Manhattan distance between T2+MPRAGE+3T runs). Surprisingly, cosine distances yielded equal sensitivity across all configurations, and the addition of the “3T” flag contributed most to the increase in sensitivity, in contrast with it’s relatively small effect on the estimated features.

### 3.3. Prediction of Participant Traits

#### 3.3.1. Sex

Classification performance was tested for three models; logistic regression (LR), support vector classifier (SVC), and random forest classifier (RFC). The default configuration scored highest both in terms of the distribution of the mean validation scores and test scores. While the complete FLAIR configuration scored higher on the test set, the results indicate that the T2w generally perform slightly better. Within the tested hyperparameter spaces, LR provided the best results across all configurations, followed by SVC and finally RFC.

#### 3.3.2. Age

Regression was tests for two models; elastic net and support vector regression (SVR). Overall, the complete FLAIR configuration yielded the highest mean validation scores, with the T2w-only configuration following closely. Elastic net generally performed slightly better than SVR within the tested hyperparameter spaces, with an average test set score close to 0.5.

## 4. Discussion

MRI preprocessing and general modeling preparation pipelines are subject to an immense number of degrees of freedom and are rarely standardized across research labs and public datasets. While some tools have managed to reach relatively widespread prevalence for particular modalities (e.g., FreeSurfer for sMRI and fMRIPrep for fMRI), these tools are highly configurable, and optimal configurations may vary significantly across acquisition protocols. In addition, some preprocessing pipelines (including but not limited to FreeSurfer) use more than one scan (e.g., in many preprocessing pipelines, an anatomical scan in required in addition to the principal modality). Again, this introduces more variance to the analysis and any consecutive modeling efforts.

In this article, we explored the differences in the structural metrics estimated by FreeSurfer with enhanced pial estimation (using T2w or FLAIR acquisitions) and explicit “3T” and “MPRAGE” flags relative to the default configuration. The estimated values were compared in three aspects: distribution of the estimated values (between and across cortical regions), within-subject similarity, and modeling performance using participant sex and age as targets. To minimize any unwanted sources of variance, all scans were acquired using a single MRI scanning protocol conducted on a single scanner and preprocessed on the same computer with the latest version of FreeSurfer available at the time (v7.2.0).

Exploration of the estimated cortical thickness across configurations revealed that using T2w-assissted workflows displays a tendency to increase the estimated values globally, whereas FLAIR shows a general decrease, which is strongly expressed in gyri and often contradicted in sulci. These results are surprising, considering these parameters are mutually exclusive and both meant to improve pial surface estimation. Cortical thickness is calculated by estimating the distance between the pial surface and the white matter surface, and therefore an improvement in pial surface estimation should improve cortical thickness estimation. However, region-wise trends would be expected to be largely similar across methods. By displaying opposite differences in the estimated cortical thickness, T2w and FLAIR configurations can’t possibly both be considered to improve pial surface estimation. While they both have a significant effect, only one could be closer to the ground truth.

In addition to the contrary trends displayed for the average, the standard deviation of the cortical thickness was found to decrease significantly in both methods. Similarly to the difference in the estimated average, this trend was relatively uniform for T2w configurations in comparison with the results using FLAIR. Unlike the average, differences in the estimated standard deviation showed a stronger regional effect, with opposite effects for neighboring regions such as the inferior temporal gyrus and the temporal pole, the medial and lateral occipitotemporal gyri, and more. Finally, FLAIR configurations also showed a general decrease in the estimated gray matter volume, which was most strongly expressed in the frontal lobe, and particularly in the medial and superior frontal gyri. Accounting for these differences is a fundamental step towards being able to successfully harmonize public datasets and significantly increase the power of derived statistical procedures.

While the accuracy of the estimated metrics in relation to the ground truth is unknown, using similarity learning and exploring the performance of various models fitted with the generated brain features provides some insight into how informative they are. Assuming brains normally maintain much greater similarity to themselves over any other brain at any time point, the calculated similarity metrics serve as an estimate of each method’s reliability. All configurations showed very high (>94.4%) sensitivity to within-subject scans using simple rule-based classification based on the minimal between-subject distance. The best results were given by calculating Manhattan distances for the “complete” T2w configuration (i.e., T2+MPRAGE+3T), with 99.3% sensitivity. Overall, Manhattan distances yielded the highest scores and cosine distance showed the most stable results across configurations. Interestingly, contrary to its relatively small effect on the estimated anatomical features, the “3T” had the largest contribution to the increase in sensitivity scores.

In terms of modeling performance by execution configuration, the results reveal significant differences with target-specific trends. Sex prediction showed the best results using the default configuration, followed by T2w and then FLAIR configurations, whereas age prediction scored highest with FLAIR configurations, followed by the T2w-only configuration, and only then the default. The reasons for these particular differences might make an interesting topic for discussion and perhaps further research. More importantly, these results suggest that preprocessing configurations should be tuned similarly to model parameters to optimize for target-specific modeling tasks. While theoretically appealing, implementing this at scale raises substantial computational challenges. To be able to fully utilize the valuable neuroimaging datasets which are continuously being curated to research finer effects, resources will need to be pooled, and new distributed research data management solutions will need to be adopted globally.

Another effect of the increase in sample size is that the distribution of data draws further away from their raw form. For this reason, it is crucial we develop heuristics which will facilitate highly powered hypothesis testing and reproducible results over time. Efforts to combine public datasets are prone to technical challenges, which may function as an “entry barrier” for participation. In addition to those, there are also imminent statistical challenges that may not be fully addressed and adequately mediated for many years before theoretical and practical issues are raised. Anticipating and preventing as many of these challenges as possible constitutes an infrastructural investment that could make a significant difference in the success and long-term reliability of future academic endeavors in the field.

In this article, we complemented prior investigations into sources of variance in FreeSurfer results by exploring the effects of a selected subset of executions configurations on the estimated structural metrics. In addition, these results were compared in terms of age and sex modeling performance using a predetermined sample of estimators and hyperparameter grids. Our results emphasize the significance of easily overlooked differences in preprocessing parameters on fundamental feature extraction results, and the unforeseeable effects these may have on consecutive modeling efforts.

---

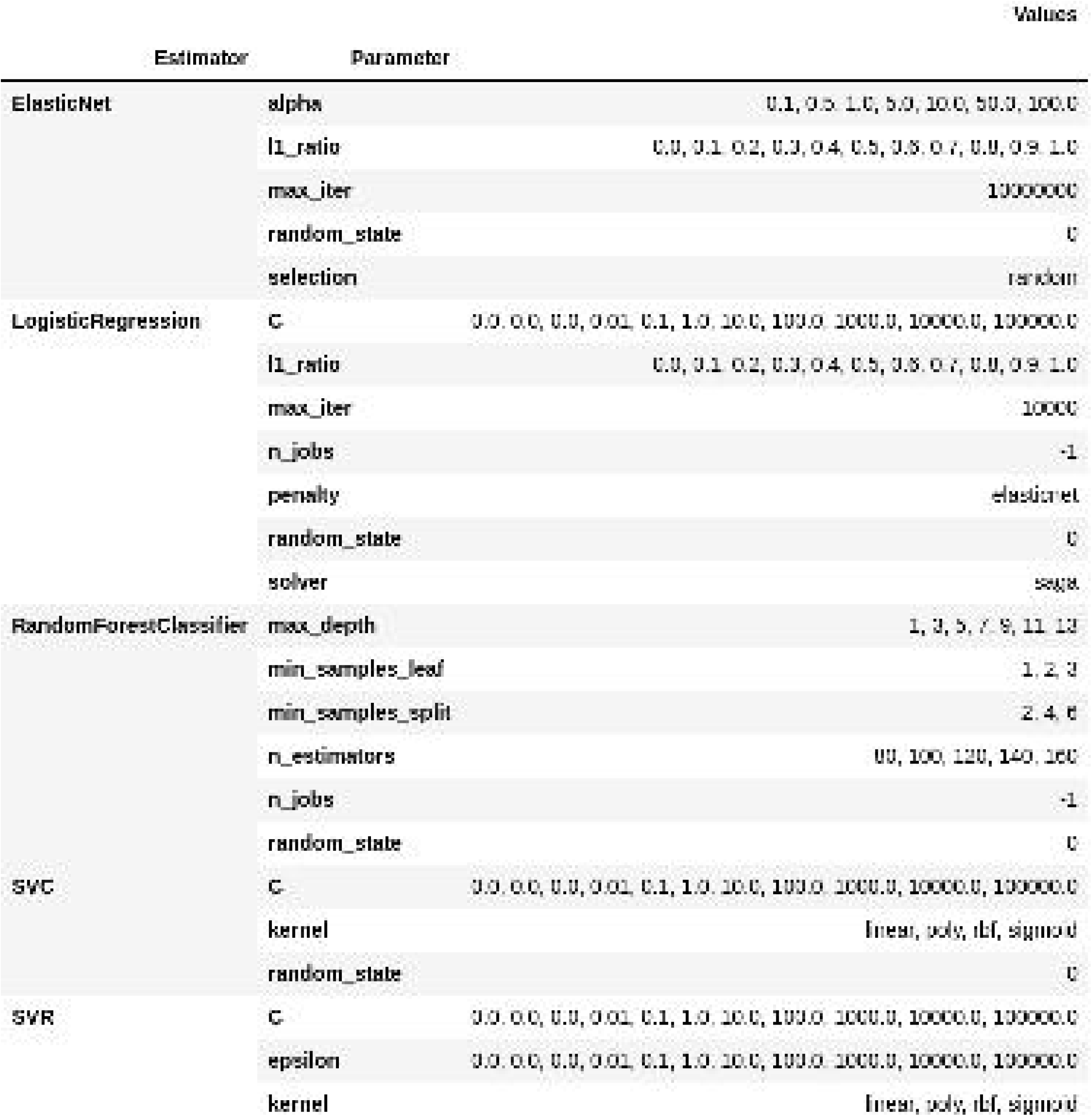

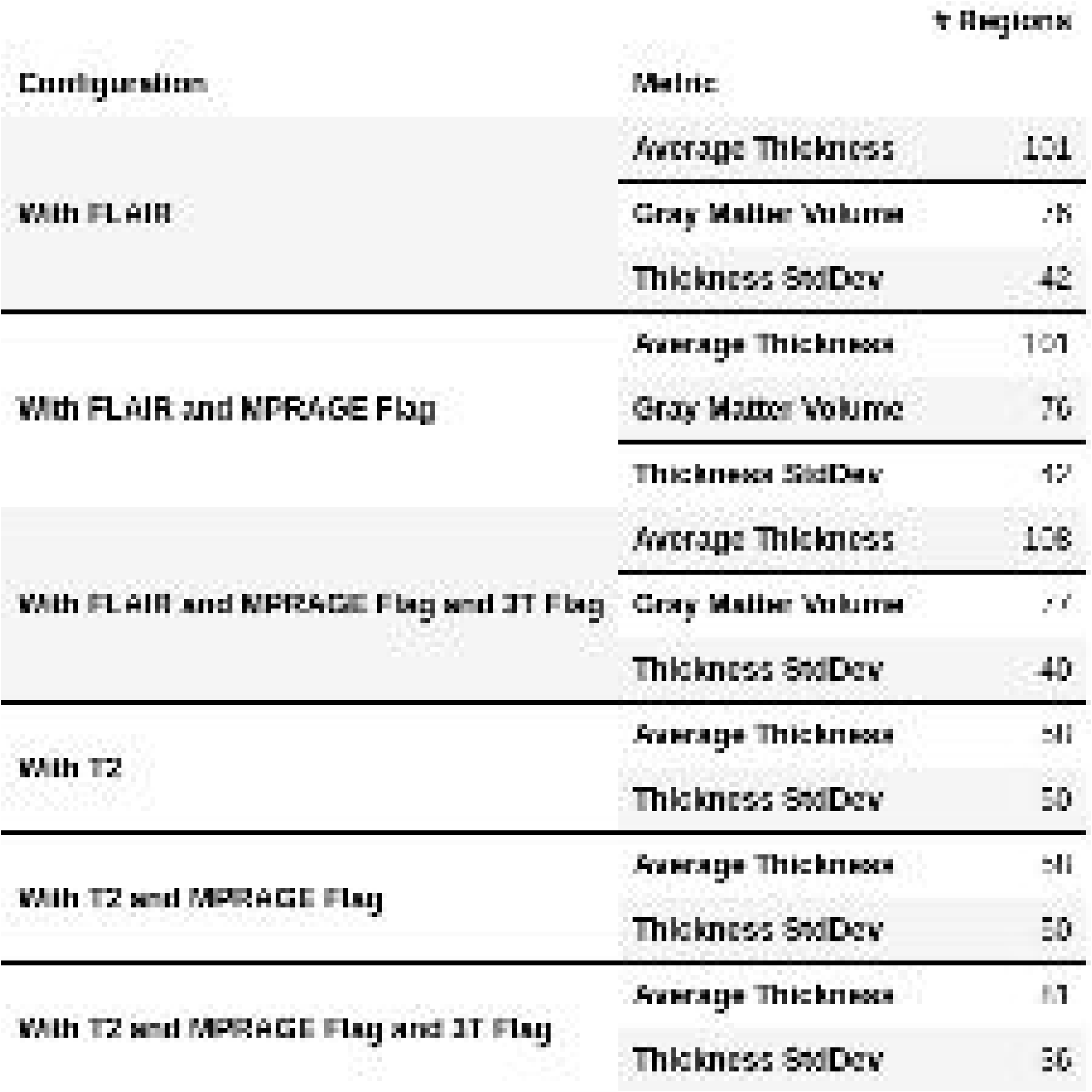

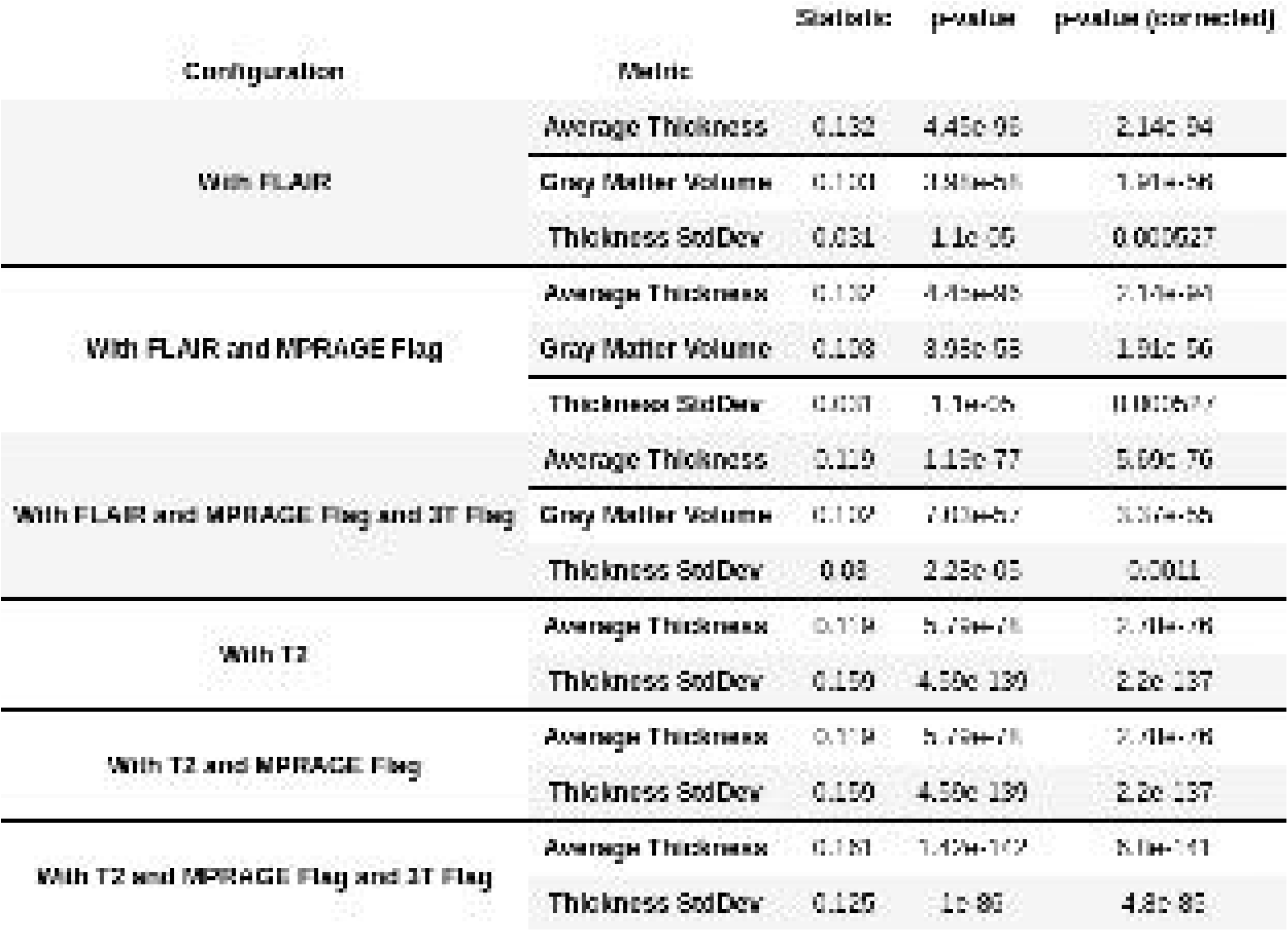

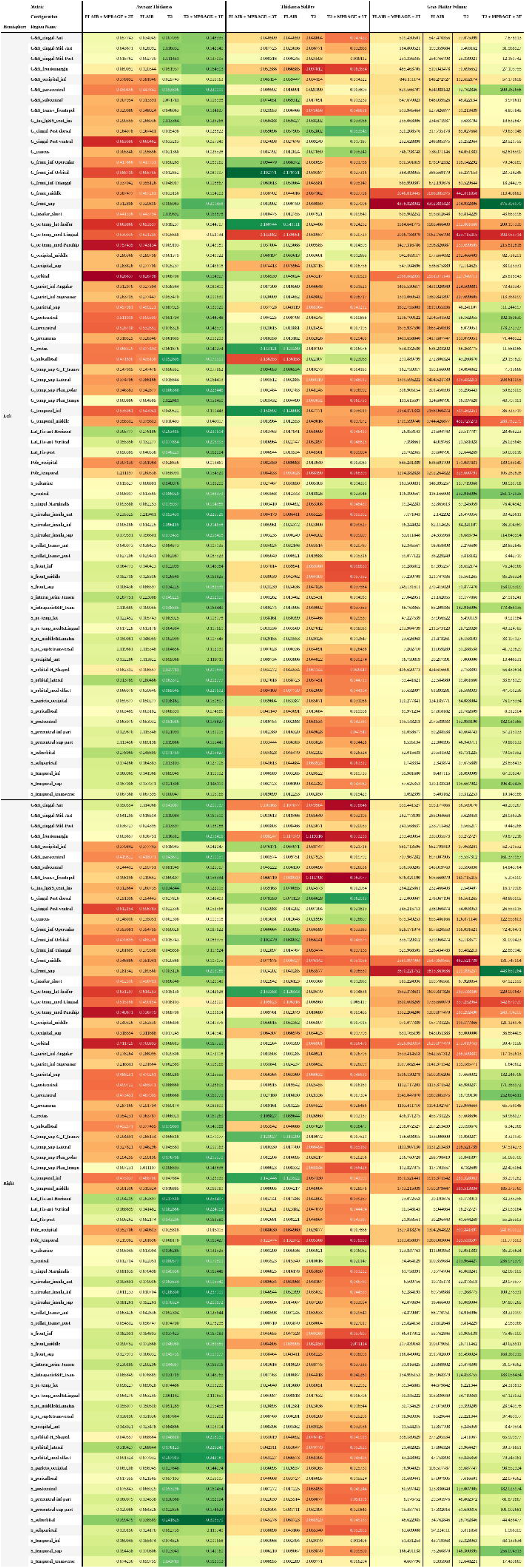

## Bibliography

Aggarwal, C. C., Hinneburg, A., & Keim, D. A. (2001). On the Surprising Behavior of Distance Metrics in High Dimensional Space. In J. Van den Bussche & V. Vianu (Eds.), Database Theory—ICDT 2001 (Vol. 1973, pp. 420–434). Springer Berlin Heidelberg. https://doi.org/10.1007/3-540-44503-X_27

Bonferroni, C. E. (1935). Il calcolo delle assicurazioni su gruppi di teste. Studi in Onore Del Professore Salvatore Ortu Carboni, 13–60.

Borghi, J. A., & Gulick, A. E. V. (2018). Data management and sharing in neuroimaging: Practices and perceptions of MRI researchers. PLOS ONE, 13(7), e0200562. https://doi.org/10.1371/journal.pone.0200562

Botvinik-Nezer, R., Holzmeister, F., Camerer, C. F., Dreber, A., Huber, J., Johannesson, M., Kirchler, M., Iwanir, R., Mumford, J. A., Adcock, R. A., Avesani, P., Baczkowski, B. M., Bajracharya, A., Bakst, L., Ball, S., Barilari, M., Bault, N., Beaton, D., Beitner, J., … Schonberg, T. (2020). Variability in the analysis of a single neuroimaging dataset by many teams. Nature, 582(7810), 84–88. https://doi.org/10.1038/s41586-020-2314-9

Dickerson, B. C., Fenstermacher, E., Salat, D. H., Wolk, D. A., Maguire, R. P., Desikan, R., Pacheco, J., Quinn, B. T., Van der Kouwe, A., Greve, D. N., Blacker, D., Albert, M. S., Killiany, R. J., & Fischl, B. (2008). Detection of cortical thickness correlates of cognitive performance: Reliability across MRI scan sessions, scanners, and field strengths. NeuroImage, 39(1), 10–18. https://doi.org/10.1016/j.neuroimage.2007.08.042

Fischl, B. (2012). FreeSurfer. NeuroImage, 62(2), 774–781. https://doi.org/10.1016/j.neuroimage.2012.01.021

Fischl, B., van der Kouwe, A., Destrieux, C., Halgren, E., Ségonne, F., Salat, D. H., Busa, E., Seidman, L. J., Goldstein, J., Kennedy, D., Caviness, V., Makris, N., Rosen, B., & Dale, A. M. (2004). Automatically parcellating the human cerebral cortex. Cerebral Cortex, 14(1), 11–22. https://doi.org/10.1093/cercor/bhg087

Gronenschild, E. H. B. M., Habets, P., Jacobs, H. I. L., Mengelers, R., Rozendaal, N., Os, J. van, & Marcelis, M. (2012). The Effects of FreeSurfer Version, Workstation Type, and Macintosh Operating System Version on Anatomical Volume and Cortical Thickness Measurements. PLOS ONE, 7(6), e38234. https://doi.org/10.1371/journal.pone.0038234

Han, X., Jovicich, J., Salat, D., van der Kouwe, A., Quinn, B., Czanner, S., Busa, E., Pacheco, J., Albert, M., Killiany, R., Maguire, P., Rosas, D., Makris, N., Dale, A., Dickerson, B., & Fischl, B. (2006). Reliability of MRI-derived measurements of human cerebral cortical thickness: The effects of field strength, scanner upgrade and manufacturer. NeuroImage, 32(1), 180–194. https://doi.org/10.1016/j.neuroimage.2006.02.051

Hedges, E. P., Dimitrov, M., Zahid, U., Brito Vega, B., Si, S., Dickson, H., McGuire, P., Williams, S., Barker, G. J., & Kempton, M. J. (2022). Reliability of structural MRI measurements: The effects of scan session, head tilt, inter-scan interval, acquisition sequence, FreeSurfer version and processing stream. NeuroImage, 246, 118751. https://doi.org/10.1016/j.neuroimage.2021.118751

Horien, C., Noble, S., Greene, A. S., Lee, K., Barron, D. S., Gao, S., O’Connor, D., Salehi, M., Dadashkarimi, J., Shen, X., Lake, E. M. R., Constable, R. T., & Scheinost, D. (2021). A hitchhiker’s guide to working with large, open-source neuroimaging datasets. Nature Human Behaviour, 5(2), 185–193. https://doi.org/10.1038/s41562-020-01005-4

Jovicich, J., Czanner, S., Han, X., Salat, D., van der Kouwe, A., Quinn, B., Pacheco, J., Albert, M., Killiany, R., & Blacker, D. (2009). MRI-derived measurements of human subcortical, ventricular and intracranial brain volumes: Reliability effects of scan sessions, acquisition sequences, data analyses, scanner upgrade, scanner vendors and field strengths. NeuroImage, 46(1), 177–192. https://doi.org/10.1016/j.neuroimage.2009.02.010

Knussmann, G. N., Anderson, J. S., Prigge, M. B. D., Dean, D. C., Lange, N., Bigler, E. D., Alexander, A. L., Lainhart, J. E., Zielinski, B. A., & King, J. B. (2022). Test-retest reliability of FreeSurfer-derived volume, area and cortical thickness from MPRAGE and MP2RAGE brain MRI images. Neuroimage: Reports, 2(2), 100086. https://doi.org/10.1016/j.ynirp.2022.100086

Laird, A. R. (2021). Large, open datasets for human connectomics research: Considerations for reproducible and responsible data use. NeuroImage, 244, 118579. https://doi.org/10.1016/j.neuroimage.2021.118579

Lehmann, M., Douiri, A., Kim, L. G., Modat, M., Chan, D., Ourselin, S., Barnes, J., & Fox, N. C. (2010). Atrophy patterns in Alzheimer’s disease and semantic dementia: A comparison of FreeSurfer and manual volumetric measurements. NeuroImage, 49(3), 2264–2274. https://doi.org/10.1016/j.neuroimage.2009.10.056

Salat, D., Greve, D., Pacheco, J., Quinn, B., Helmer, K., Buckner, R., & Fischl, B. (2009). Regional white matter volume differences in nondemented aging and Alzheimer’s disease. NeuroImage, 44(4), 1247–1258. https://doi.org/10.1016/j.neuroimage.2008.10.030

Seabold, S., & Perktold, J. (2010). Statsmodels: Econometric and Statistical Modeling with Python. 92–96. https://doi.org/10.25080/Majora-92bf1922-011

Valizadeh, S. A., Liem, F., Mérillat, S., Hänggi, J., & Jäncke, L. (2018). Identification of individual subjects on the basis of their brain anatomical features. Scientific Reports, 8(1), Article 1. https://doi.org/10.1038/s41598-018-23696-6

Van Horn, J. D., & Toga, A. W. (2009). Is it time to re-prioritize neuroimaging databases and digital repositories? NeuroImage, 47(4), 1720–1734. https://doi.org/10.1016/j.neuroimage.2009.03.086

Virtanen, P., Gommers, R., Oliphant, T. E., Haberland, M., Reddy, T., Cournapeau, D., Burovski, E., Peterson, P., Weckesser, W., Bright, J., van der Walt, S. J., Brett, M., Wilson, J., Millman, K. J., Mayorov, N., Nelson, A. R. J., Jones, E., Kern, R., Larson, E., … van Mulbregt, P. (2020). SciPy 1.0: Fundamental algorithms for scientific computing in Python. Nature Methods, 17(3), Article 3. https://doi.org/10.1038/s41592-019-0686-2

